# Analysis of gastric cancer transcriptome allows the identification of histotype specific molecular signatures with prognostic potential

**DOI:** 10.1101/2021.02.02.429357

**Authors:** Adriana Carino, Luigina Graziosi, Silvia Marchianò, Michele Biagioli, Elisabetta Marino, Valentina Sepe, Angela Zampella, Eleonora Distrutti, Annibale Donini, Stefano Fiorucci

**Affiliations:** University of Perugia, Department of Medicine and Surgery, Perugia, Italy; Azienda Ospedaliera di Perugia, Perugia, Italy; University of Naples Federico II, Department of Pharmacy, Naples, Italy

**Keywords:** gastric cancer, adenocarcinoma, biomarkers, tumor microenvironment, transcriptome expression

## Abstract

Gastric cancer is the fifth most common malignancy but the third leading cause of cancer-associated mortality worldwide. Therapy for gastric cancer remain largely suboptimal making the identification of novel therapeutic targets an urgent medical need. In the present study we have carried out a high-throughput sequencing of transcriptome expression in patients with gastric cancers. Twenty-four patients, among a series of 53, who underwent an attempt of curative surgery for gastric cancers in a single center, were enrolled. Patients were sub-grouped according to their histopathology into diffuse and intestinal types, and the transcriptome of the two subgroups assessed by RNAseq analysis and compared to the normal gastric mucosa. The results of this investigation demonstrated that the two histopathology phenotypes express two different patterns of gene expression. A total of 2064 transcripts were differentially expressed between neoplastic and non neoplastic tissues: 772 were specific for the intestinal type and 407 for the diffuse type. Only 885 transcripts were simultaneously differentially expressed by both tumors. The per pathway analysis demonstrated an enrichment of extracellular matrix and immune dysfunction in the intestinal type including CXCR2, CXCR1, FPR2, CARD14, EFNA2, AQ9, TRIP13, KLK11 and GHRL. At the univariate analysis reduced levels AQP9 was found to be a negative predictor of 4 years survival. In the diffuse type low levels CXCR2 and high levels of CARD14 mRNA were negative predictors of 4 years survival. In summary, we have identified a group of genes differentially regulated in the intestinal and diffuse histo-types of gastric cancers with AQP9, CARD14 and CXCR2 impacting on patients prognosis, although CXCR2 is the only factor independently impacting overall survival.

**Simple summary:** Gastric cancer is the fifth most common malignancy and the third leading cause of cancer-associated mortality worldwide. Although several new pharmacological approaches are currently developed, surgery remains the unique valid option of treatment but survival remains very poor over the last decades. Therefore, understanding the underlying molecular mechanisms of the gastric carcinogenesis and identifying sensitive biomarkers could be helpful for the prevention and treatment of the disease. Currently, the high-throughput sequencing techniques, in particular the transcriptomic analysis (RNA-seq) represents a validated technique to obtain a molecular characterization of human cancers. Moreover, it has been established that genetic susceptibility and environmental factors, such as microbial infections may contribute to carcinogenesis. We have characterized the different patterns of gene expression, using RNA-seq analysis and correlated these findings with gastric cancer histological subtypes.

## Introduction

Gastric cancer is a highly prevalent cancer representing the fifth most frequent cancer worldwide [1–3]. While gastric cancer incidence has shown a trend of reduction over the last decades, gastric cancer-related mortality remains the third cause of cancer-related deaths worldwide. In Western countries, due to a lack of validated cancer screening programs, the large majority patients with gastric cancer are diagnosed in the advanced stages, greatly limiting the therapeutic options. Surgery remains the only potentially curative treatment, although current evidence supports adoption of perioperative therapies to improve a patient’s survival [2]. Peritoneal metastases are the most frequent metastases detected in patients with advanced gastric cancers and the peritoneal cavity is a common site for gastric cancer recurrence following surgery [4] [5]. Overall, the presence of peritoneal carcinosis is associated to reduced survival rates and overwhelming symptoms [6] [7].

Two histopathology subtypes of gastric adenocarcinomas, intestinal (well-differentiated) and diffuse (undifferentiated), with a distinct morphologic appearance, pathogenesis, and genetic profiles have been identified [8–10]. The diffuse gastric cancer type is clinically more aggressive and associates with a higher rate of peritoneal involvement compared to the intestinal type [11,12]. However, the current histopathologic system fails to reflect the molecular and genetic heterogeneity of gastric cancers, and it is of clinical relevance to molecularly investigate gastric cancers in the attempt to identify novel targets for the prevention and treatment [9].

Several studies have shown a robust molecular heterogeneity of gastric adeno-carcinomas leading to different molecular classifications [11,13,14]. A widely used molecular classification proposed by the Genome Atlas Research Network Group (TCGA) has identified four major tumor molecular subtypes: Epstein–Barr virus positive, microsatellite unstable tumors, genomically stable tumors and tumors with chromosomal instability [14]. Although these subtypes have shown poor correlation with the prognosis, they have proven partially helpful in the selection of chemotherapy approaches suggesting that molecular profiling rather than histology could be implemented as a guide for the choice of treatment modality. Currently, only few biomarkers are available to predict treatment effectiveness in gastric cancer’s patients including the level of expression of human epidermal growth factor receptor 2 (HER2) for trastuzumab and the programmed death-ligand 1 (PD-L1) for pembrolizumab [15,16], the last one allowed as a second line therapy for metastatic disease.

The introduction of Next Generation Sequencing (NGS), such as the RNA-sequencing technology, allows the application of large-scale functional genomics to cancer research and its application in clinical settings might allow the identification of individual gene expression profiles to be used as a potential biomarkers in the treatment of gastric cancer [17,18]. Previous studies have investigated differentially expressed genes between gastric cancer tissues and healthy gastric mucosa [17,19,20]. Moreover, several studies have identified stage-specific gene expression profiles and histological specific-gene profiles [21,22]. In this study, we report an integrative analyses of transcriptome sequencing (RNA-seq) and clinical and pathological characterization of patients with gastric cancers. We performed the NGS analysis based on histological types. Our principal goal was to identify new transcripts to be used as gastric cancer biomarkers and to which attribute a prognostic survival value. This novel comparative analysis generated a large amount of information that could be exploited to identify underlying molecular mechanisms of carcinogenesis, detection of disease markers and the identification of novel therapeutic targets.

## Materials and Methods

### Patients and specimens

Surgical specimen of 53 patients curatively operated for gastric cancers, were collected and analyzed according to the Helsinky declaration. Permission to collect post-surgical samples was granted to Prof. Fiorucci by the ethical committee of Umbria (CEAS). Permit FI00001, n. 2266/2014 granted on February 19, 2014. An informed written consent was obtained by each patient before surgery.

Clinical data of these patients were retrieved from a prospectively collected database of GC patients operated between October 2014 and August 2017 at the Department of Surgery, Santa Maria della Misericordia Hospital (Italy). None of the patients received chemotherapy or radiation before surgery. All patients were followed up regularly until death every 6 months for the first 2 years from surgery and every year thereafter.

The study population includes 24 patients that were selected according to preoperative factors such as sex, age, gender, preoperative serum albumin level, preoperative N/l ratio (neutrophils to lymphocytes ratio); surgery related factors such as surgery type, lymphadenectomy level dissection, number of lymph nodes retrieved; associated organ resections and tumor related factors such as Lauren’s Hystotype, Tumor location, Pathological stage according to AJCC tnm 8^th^ editon [23] and peritoneal carcinomatosis development.

Venous blood sample was taken either the day before surgery or few days immediately before and collected in ethylenediaminetetraacetic acid-containing tube. The normal range of white blood cell (WBC) count was from 4,000 to 10,800 cells/mm3 and 3.5-4. The normal range of albumin was from 3,5 g/dl to 5,2 g/dl. N/L was calculated as neutrophil count divided by lymphocyte count. The patients were dichotomized at the median value of NLR, whereas patients were dichotomized according to the lower physiologic albumin levels for both intestinal and diffuse groups (Table 1).

**Table 1.** Clinical and pathological characteristics of the study population (n=24).

| Patients |  | Intestinal (n=12) | Diffuse (n=12) | P |
| --- | --- | --- | --- | --- |
| Age* |  | 72.2 ±5.6 | 71.7±12.5 | N.S. |
| Gender: |  |  |  |  |
|  | • M | 9 (75%) | 6 (50%) | N.S. |
|  | • F | 3 ( 25%) | 6 (50%) |  |
| pT: |  |  |  |  |
|  | • 1 | 0 | 0 | N.S. |
|  | • 2 | 0 | 1 (8.3%) |  |
|  | • 3 | 6 (50%) | 5 ( 41.7%) |  |
|  | • 4 | 6 (50%) | 6 (50%) |  |
| pN: |  |  |  |  |
|  | • No | 1 (8.3%) | 1 (8.3%) | N.S. |
|  | • N1 | 1 (8.3%) | 0 |  |
|  | • N2 | 3 (25%) | 4 (33.3%) |  |
|  | • N3 | 7 (58.4) | 7 (58.4) |  |
| Stage: |  |  |  |  |
|  | • I | 0 | 0 | N.S. |
|  | • II | 1 (8.3%) | 0 |  |
|  | • III | 8 ( 66.7%) | 6 (50%) |  |
|  | • IV | 3(25%) | 6 (50%) |  |
| Lymphonodal Harvasted**: |  | 38.5 | 37 | N.S. |
| Lymphonodal Ratio** |  | 0.2 | 0.3 | N.S. |
| Tumor Location: |  |  |  |  |
|  | • U | 2 (16.7%) | 4 (33.3%) | N.S. |
|  | • M | 6 (50%) | 5 (41.7%) |  |
|  | • L | 4 (33.3%) | 3 (25%) |  |
| Lymphoadenectomy: |  |  |  |  |
|  | • D1 | 1 (8.3%) | 2 (16.7%) | N.S. |
|  | • D2 | 11 (91.7%) | 9 ( 75%) |  |
|  | • D3 | 0 | 1 (8.3%) |  |
| Multiorgan resection: |  | 2 (16.7%) | 3 (25%) | N.S. |
| Peritoneal Carcinomatosis development: |  | 4 (33.3%) | 8 (66.6%) | N.S. |
| N/L ratio**: |  | 2.3 | 3.5 | N.S. |
| Sieric Albumin*** |  | 3.5 | 3.5 | N.S. |

Finally, patients were separated in to two subgroup according to the median value of gene expression to perform OS analysis.

### Statistical analysis

Patients descriptive analysis was generated, and their differences were investigated using Student t-test for quantitative data; for qualitative data, we used either Fisher’s exact test or chi-square test. To compare overall survival (OS) between groups, the cumulative survival proportions were calculated using the product limit method of Kaplan-Meier, and differences were evaluated using the log-rank test. Only variables that achieved statistical significance in the univariate analysis were subsequently evaluated in the multivariate analysis using Cox’s proportional hazard regression model. A p value of less than 0.05 was considered statistically significant. All statistical analyses were performed using the MedCalc Statistical Software version 14.8.1 (MedCalc Software bvba, Ostend, Belgium) and PRISM 7.2 Graph PAD.

### AmpliSeq Transcriptome

High-quality RNA was extracted from tumor gastric mucosa and healthy mucosa using the PureLink^™^ RNA Mini Kit (Thermo Fisher Scientific), according to the manufacturer’s instructions. RNA quality and quantity were assessed with the Qubit^®^ RNA HS Assay Kit and a Qubit 3.0 fluorometer followed by agarose gel electrophoresis. Libraries were generated using the Ion AmpliSeq^™^ Transcriptome Human Gene Expression Core Panel and Chef-Ready Kit (Thermo Fisher Scientific), according the manufacturer’s instructions. Briefly, 10 ng of RNA was reverse transcribed with SuperScript^™^ Vilo^™^ cDNA Synthesis Kit (Thermo Fisher Scientific, Waltham, MA) before library preparation on the Ion Chef^™^ instrument (Thermo Fisher Scientific, Waltham, MA). The resulting cDNA was amplified to prepare barcoded libraries using the Ion Code™ PCR Plate, and the Ion AmpliSeq^™^ Transcriptome Mouse Gene Expression Core Panel (Thermo Fisher Scientific, Waltham, MA), Chef-Ready Kit, according to the manufacturer’s instructions. Barcoded libraries were combined to a final concentration of 100 pM, and used to prepare Template-Positive Ion Sphere^™^ (Thermo Fisher Scientific, Waltham, MA) Particles to load on Ion 540^™^ Chips, using the Ion 540^™^ Kit-Chef (Thermo Fisher Scientific, Waltham, MA). Sequencing was performed on an Ion S5^™^ Sequencer with Torrent Suite^™^ Software v6 (Thermo Fisher Scientific). The analyses were performed with a range of fold <−2 and >+2 and a p value <0.05, using Transcriptome Analysis Console Software (version 4.0.2), certified for AmpliSeq analysis (Thermo-Fisher). The transcriptomic data have been deposited as dataset on Mendeley data repository (10.17632/d3ykf83tyv.1).

### Functional enrichment analysis

DAVID software was employed to identify significantly enriched Gene Ontology functions in biological processes (BP), molecular function, and cellular component (CC) categories for histotype-specific genes [24].

## Results

### Patients

This study includes 24 gastric cancer patients who underwent resection surgery in our department between October 2014 to August 2017. This cohort of patients was further subdivided into two groups according to Lauren classification, as such two group of gastric cancer patients were identified: “intestinal group” and “diffuse group” (Table 1). The two groups include 12 patients each and were highly homogeneous as indicated in table 1, in term of mean age, gender and cancer stage. All patients underwent either a total or subtotal gastrectomy plus D1, D2 or D3 lymphadenectomy with a median number of harvested lymph nodes of 38.5 for intestinal group and 37 for diffuse group. The distribution of cancer stages (TNM8) was as follows: stage II: 1 (8.3%), stage III: 8 (66.7%), stage IV: 3 (25%) for intestinal group; stage III: 6 (50%), stage IV: 6 (50%) for diffuse group (Table 1). For survival analysis two patients in each group were excluded due to early postoperative death.

### Transcriptome Analysis: Identification of common markers that characterized both diffuse and intestinal gastric cancer from healthy mucosa

We performed an AmpliSeq Transcriptome analysis (RNA-seq) of 24 gastric tumors (and their matched normal tissues), according to the Lauren classification, as described in Material and Method section. As shown in Figure 1, the PCoA analysis revealed that healthy samples showed a homogeneous distribution, whereas both diffuse and intestinal tumors showed dissimilarities compared to normal tissues, but their signals only partially overlap (Figure 1A). The Scatter Plots depicted in Figure 1B, identify transcripts differentially expressed between diffuse tumor and healthy mucosa, or intestinal tumor and healthy mucosa.

**Figure 1.**
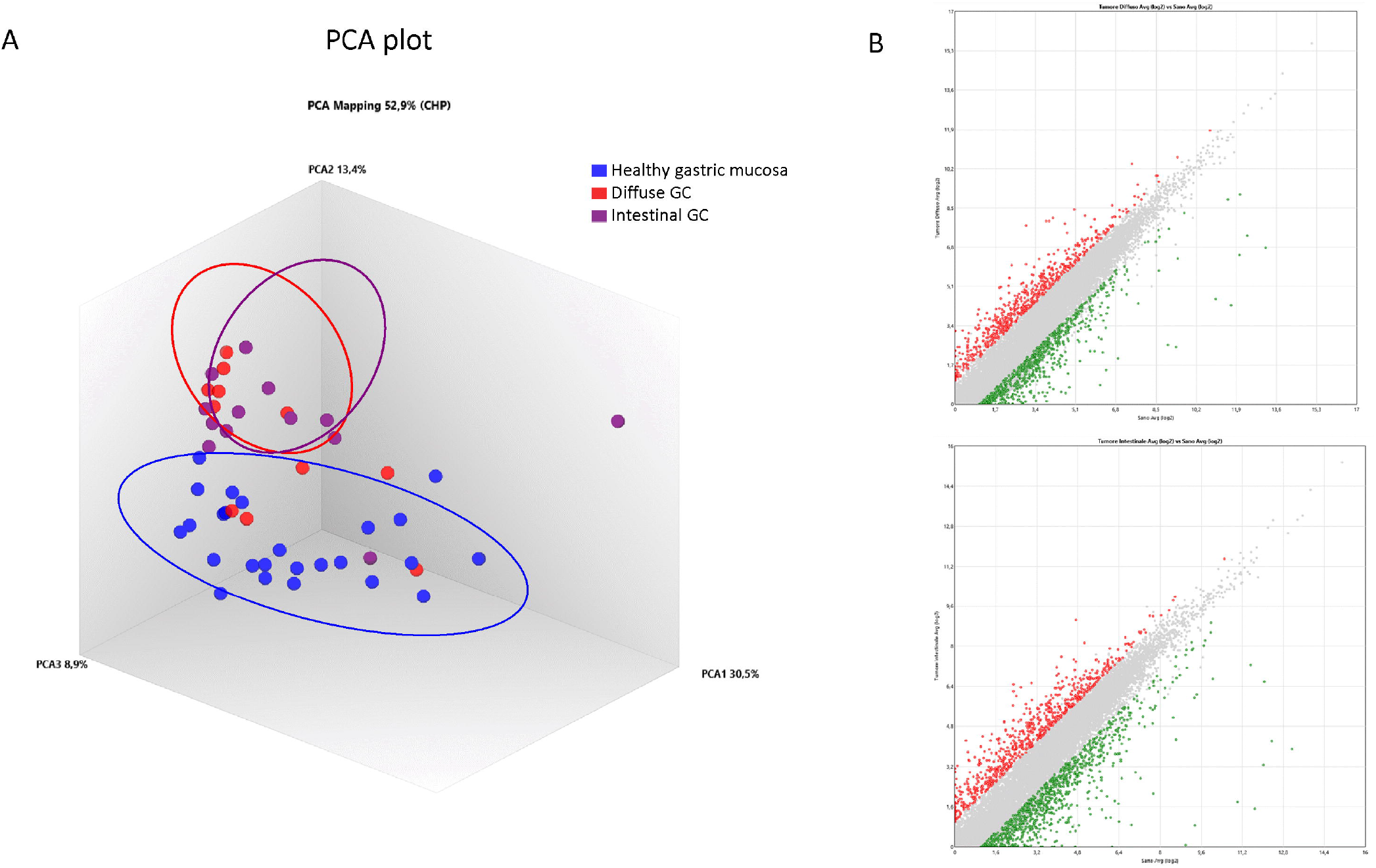
RNA sequencing of Diffuse and Intestinal Gastric Cancer. (A) Heterogeneity characterization of gastric samples showed by principal component analysis (PCA) plot. (B) Scatter plots of transcripts differentially expressed between diffuse gastric cancer and healthy mucosa or intestinal gastric cancer and healthy mucosa. (Fold Change <-2 or >+2, p value <0.05).

The Venn Diagram analysis of differentially expressed transcripts, confirmed that the signals of diffuse and intestinal gastric cancer only partially overlap. We have identified the subset AB (885 transcripts) containing genes differentially expressed both in diffuse and intestinal gastric cancer vs healthy mucosa (Figure 2). These transcripts resulted modulated in same direction in both gastric cancer histotypes, confirming the existence of common molecular mechanisms underlying the development of the two main histological types of gastric cancer.

**Figure 2.**
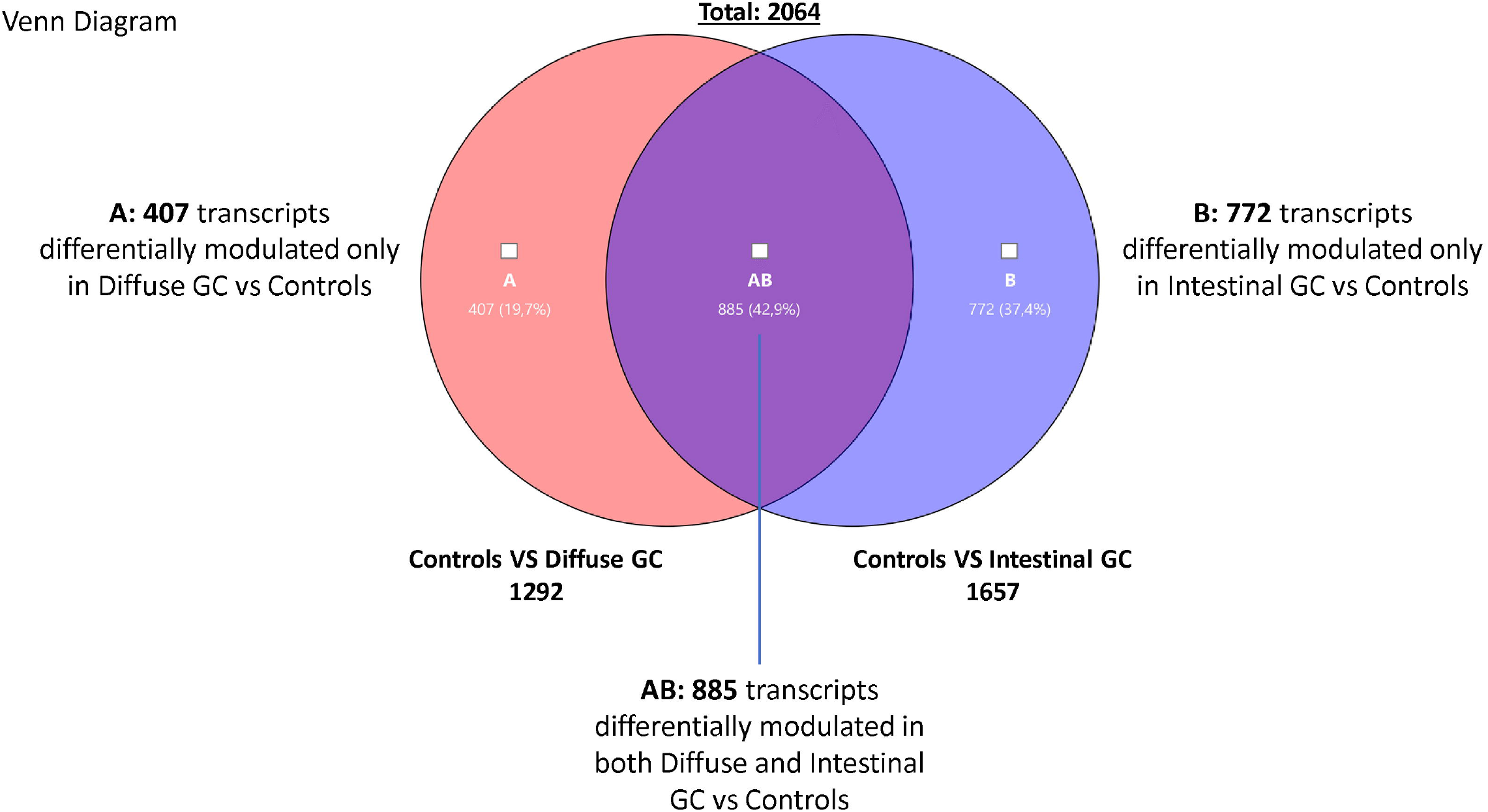
Transcriptome analysis. Venn Diagram analysis of differentially expressed genes in Diffuse (subset A) and Intestinal (subset B) gastric cancer samples compared with healthy mucosa, showing the overlapping region (identified AB subset) containing transcripts differentially expressed in both gastric cancer hystotypes (Fold Change <-2 or >+2, p value <0.05).

We have also detected transcripts differentially modulated only in diffuse gastric cancer vs healthy mucosa (Subset A containing 407 genes), and transcripts differentially modulated only in intestinal cancer vs healthy mucosa (Subset B containing 772 genes). These findings suggest that each gastric cancer histotype is characterized by specific molecular patterns. Therefore, we have performed a *per* pathways analysis of these different gene subsets using TAC software, to better dissect the most modulated mechanisms in diffuse and intestinal gastric cancers. For each subset (AB, A and B) we found several pathways that can be grouped in three principal clusters: a) proliferation, differentiation and metabolism; b) inflammation and c) signaling. As expected, we found that the two types of gastric cancer (Subset AB) showed similar features regarding the modulation of genes involved in cell cycle, mitosis, cell division, DNA replication, extracellular matrix, as well as in the regulation of inflammation (IL-18, Chemokines, Cytokines, IL6 signaling pathways), or in cancer development and progression (PI3K-Akt-mTOR, VEGFA-VEGFR2, MAPK, Ras, EGF/EGFR signaling pathways), which differentiate both histotypes from healthy mucosa (see Supplementary Tables 1, 2 and 3).

In particular, we found an upregulation of several cell cycle regulators (Supplementary Table 1) such as the cyclin-dependent kinase CDK1, that induces the growth of gastric cancer cells [25], the M-phase inducers CDC25A and CD2C5B, which increased expression represents an early event in gastric carcinogenesis common to both diffuse and intestinal cancer [26], the Cyclin B1 and B2 (CCNB1, CCNB2), upregulated to promote gastric cancer cell proliferation and tumor growth [27,28], or the E2Fs family and the correlated MYBL2 proto-oncogene, associated with cancer progression and poor overall survival [29,30]. Importantly we found also a modulation of genes involved in Epithelial to Mesenchymal Transition including TMPRSS4, that is upregulated in gastric cancer and increased the invasiveness of gastric cancer cells activating NF-Kb/MMP9 signaling [31], and Claudins (CLDN1, CLDN3, CLDN4, CLDN7), overexpressed in gastric cancer and associated with gastric cancer cell proliferation, invasion and maintenance of mesenchymal state [32–34]. Furthermore, we found an overexpression of ITGA2, which regulates Metastases and EMT [35], LAMC2 and WNT5A genes, that mediate invasion of gastric cancer cells [36]. Finally, great of relevance, we found an upregulation of Matrix Metalloproteinase family (MMP1, MMP3, MMP10, MMP12), that are overexpressed in gastric cancer as a result of NF-Kb activation thus promoting migration and invasion, and are associated with poor prognosis [37–40]. Among most downregulated genes we found GPX3, PTGER3 and LIPF (−47.5 and −435.63 of fold change for Diffuse and Intestinal tumors respectively), that resulted hypermethylated in gastric cancer [41,42]. The analysis of immune cluster revealed a great modulation of chemokine and cytokine signaling pathways (Supplementary Table 2). In particular, we found an upregulation of CCL3, CCL15, both overexpressed in the stromal compartment of gastric cancer [43], CCL20, that activates the pErk1/2-pAkt signaling via CCR6 inducing EMT pathways [44], but also CXCLs family (CXC1, CXCL2, CXCL5 and CXCL16), that are induced by Cox2/Pge2 or Wnt5a signaling [45,46], and correlate with tumor malignant progression, invasiveness, gastric cancer cell migration and metastasis via the activation of CXCR2/Stat3 pathway [47–50].

We also detected an upregulation of several cytokines including IL1β, IL11 and IL8, that promote metastasis, anti-apoptotic effects and maintenance of stemless properties by activating PI3K, Jak/Stat3 and Nf-Kb signaling pathways respectively [51–53]. Interestingly, we detected a downregulation of AQP4, which, when overexpressed, reduce gastric cancer cells proliferation [54]. The analysis of signaling cluster (Supplementary Table 3), revealed the upregulation of ANGPT2, whose overexpression promotes angiogenesis in gastric cancer [55], and S100A2, which is associated with tumor progression [56], whereas the expression of several genes was downregulated, including: CAB39L, a tumor suppressor hypermethylated in gastric cancer cells and tissues [57], HIF3A and SFRP1, targeted by several specific miRNA overexpressed in gastric cancer [58–60], and FABP4, whose expression in Tissue-resident memory T cells (Trms) infiltrating the tumor is reduced by PDL1 activation [61].

Furthermore, among overexpressed genes in both gastric cancer types, we found CEACAM1 and CEACAM6, two recognized markers of angiogenesis, invasion and metastasis in gastric cancer [62,63]. Finally, we also detected a modulation of genes that have been identified as tumor suppressors in other cancer types, including OGN, a gene that is downregulated in the breast cancer and that functions as Pi3K/Akt/mTOR suppressor [64].

### Transcriptome Analysis: Identification of markers specific for Diffuse Gastric cancer

To better characterize the specific phenotype of Diffuse Gastric cancer, we have then investigated the subset A of Venn Diagram (Figure 3). The analysis of this subset of genes, highlighted a modulation of transcripts involved in lipid metabolism and transport, metabolic pathways and transcription (Table 2).

**Figure 3.**
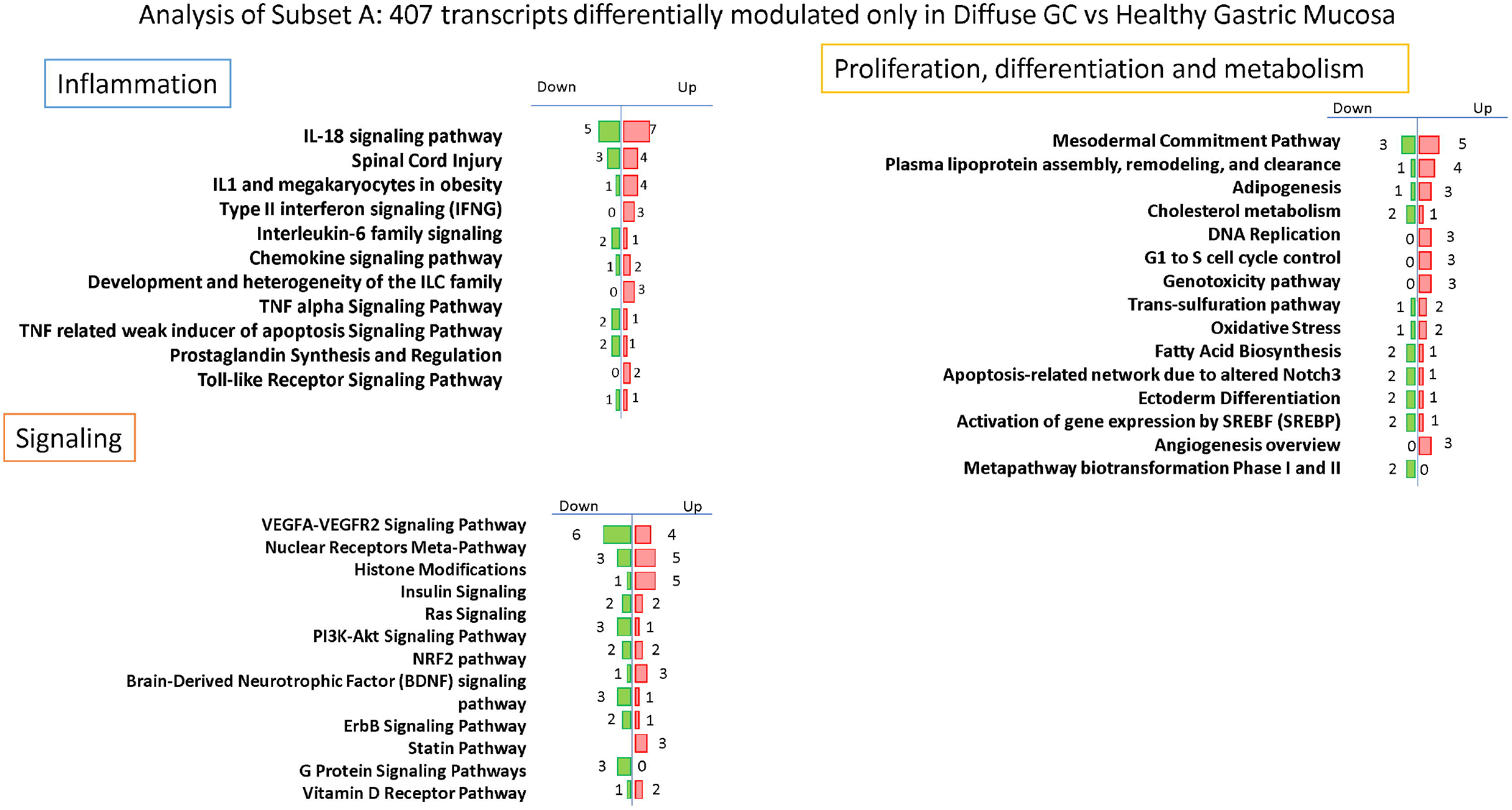
*Per* pathways analysis of Subset A. Analysis of 407 transcripts differentially modulated only in Diffuse Gastric Cancer vs Healthy Gastric Mucosa: identification of several pathways that can be grouped in three principal clusters: Proliferation, differentiation and metabolism, Inflammation and Signalling.

**Table 2.** Principal pathways of Proliferation, differentiation and metabolism Cluster for the subset A.

| Proliferation, differentiation and metabolism | Upregulated genes | Downregulated genes |
| --- | --- | --- |
| Mesodermal Commitment Pathway | KLF5, ASCC3, <u>HNF4A</u>, HPRT1, HMGA2 | ATP8B2, PBX1, ARID5B |
| Plasma lipoprotein assembly, remodeling, and clearance | <u>APOC2</u>, <u>APOE</u>, <u>APOC1</u>, NCEH1 | <u>VLDLR</u> |
| Adipogenesis | KLF5, TRIB3, LMNA | CNTFR |
| Cholesterol metabolism (includes both Bloch and Kandutsch-Russell pathways) | <u>FASN</u> | TM7SF2, <u>CYP27A1</u> |
| DNA Replication | CDC7, POLA2, POLE2 |  |
| G1 to S cell cycle control | POLE2, CREB3L1, POLA2 |  |
| Genotoxicity pathway | HIST1H2BI, HIST1H2BM, HIST1H3D |  |
| Trans-sulfuration pathway | GCLM, DNMT1 | GGT6 |
| Oxidative Stress | HMOX1, NOX4 | MT1X |
| Fatty Acid Biosynthesis | <u>FASN</u> | ECHDC2, ACACB |
| Apoptosis-related network due to altered Notch3 | <u>APOE</u> | TRAF1, NGFRAP1 |
| Ectoderm Differentiation | STC1 | <u>CCL2</u>, CTNND2 |
| G1 to S cell cycle control | POLE2, CREB3L1, POLA2 |  |
| Genotoxicity pathway | HIST1H2BI, HIST1H2BM, HIST1H3D |  |
| Trans-sulfuration pathway | GCLM, DNMT1 | GGT6 |
| Oxidative Stress | HMOX1, NOX4 | MT1X |

In particular, we found an upregulation of several genes involved in proliferation and lipid metabolism such as HNF4A, functionally required for the development of gastric cancer regulating IDH1 [65], APOC1, recognized as new diagnostic and prognostic marker of gastric cancer [66], APOE, which is highly expressed in gastric cancers and correlates with progression and invasion [67,68], FASN, associated with diffuse gastric cancer and poor prognosis [69]. Conversely, we found a downregulation of VLDLR, whose genetic or epigenetic silencing contributes to gastric carcinogenesis [70], CYP27A1, which induces T cell dysfunction, thus promoting breast cancer progression [71] and CCL2, downregulated in diffuse gastric cancer primarily in advanced stages [72].

Although to a lesser extent, we also found a modulation of genes involved in inflammatory and signaling pathways (Supplementary Table 4 and 5). Interestingly, we found an upregulation of inflammatory genes including IL-18, marker of TAMs (Tumor associated macrophages) probably correlated with tumor invasion ability [73], IFNγ, that promotes gastric tumorigenesis in mice and regulates the expression of PD1 via JACK/Stat signaling [74,75], CCL22, that promotes EMT activating PI3K/Akt pathway and is correlated with peritoneal metastasis in gastric cancer patients [76,77], and CCL28, molecular target of Wnt/β-cathenin overexpressed in gastric cancer [78]. Furthermore, we found an overexpression of MMP-9, that promotes tumor invasion and is associated with poor prognosis in gastric cancer [79], ANXA1 and ANXA4, overexpressed in gastric cancer and associated with proliferation [80,81], AREG, that promotes malignant progression in several types of cancer [85]. Among the downregulated genes we found ADAMTS1, a metalloprotease with anti-angiogenic activity expressed at low levels in primary tumors [82]. Moreover, the 3 most overexpressed specific genes in diffuse gastric cancer resulted REG4, LCN2 and CEACAM5 (fold change of 16.97, 7.63 and 6.34 respectively): REG4 is generally overexpressed in gastric cancer and promotes peritoneal metastasis, increasing adhesion ability of gastric cancer cells [83]; LCN2 is overexpressed in gastric cancer mucosa infected with *H. pylori* and correlated with invasion, metastasis and poor prognosis of gastric cancer [84,85]; the expression of CEACAM5 is associated with tumor progression, invasion and migration and it is considered an independent prognostic predictor in patients with advanced stages of gastric cancer [86,87]. Conversely the most downregulated gene in diffuse gastric cancer subset is REG1A (fold change of −7.47), a tumor suppressor that, when overexpressed, reduces invasion and promotes apoptosis of gastric cancer cells, and that is typically downregulated in gastric cancer patients [88,89].

### Transcriptome Analysis: Identification of markers specific for Intestinal Gastric cancer

The analysis of intestinal gastric cancer subset indicated as subset B (Figure 4), immediately showed a stronger inflammatory component, likely due to its greater association with *H. pylori* infection. As shown in Table 3, the *per pathway* analysis of inflammatory cluster indicates a great modulation of genes involved in chemotaxis, inflammation, Innate and adaptative immunity (Table 3). First, we found a strong modulation of IL-18 signaling pathway with an overexpression of several genes including CCL4, whose increased expression in stromal compartment is associated with Intestinal gastric cancer [43], FN1, considered a prognostic biomarker in gastric cancer associated with a poor prognosis [90], and PTGS2, encoding for COX2 gene with a key role in the generation of the inflammatory microenvironment in tumor tissues inducing the expression of several cytokines and chemokines, which play tumor-promoting role [45,91]. In this pathway, we have also found a downregulation of CCL19, a tumor suppressor that reduces proliferation, migration and invasion in gastric cancer [92], and NR0B2, whose downregulation in renal carcinoma is associated with development and progression of cancer [93]. Moreover the, Intestinal gastric cancer group is characterized by a modulation of chemokine signaling pathway, in which we found an increased expression of CXCR2, the IL8 receptor, associated with poor prognosis and metastasis [94], and conversely a downregulation of CCL19, that suppresses proliferation, migration and invasion of gastric cancer cells [92], and CXCL14, whose promoter hypermethylation is associated with depth of penetration and prognosis of gastric cancer [95].

**Figure 4.**
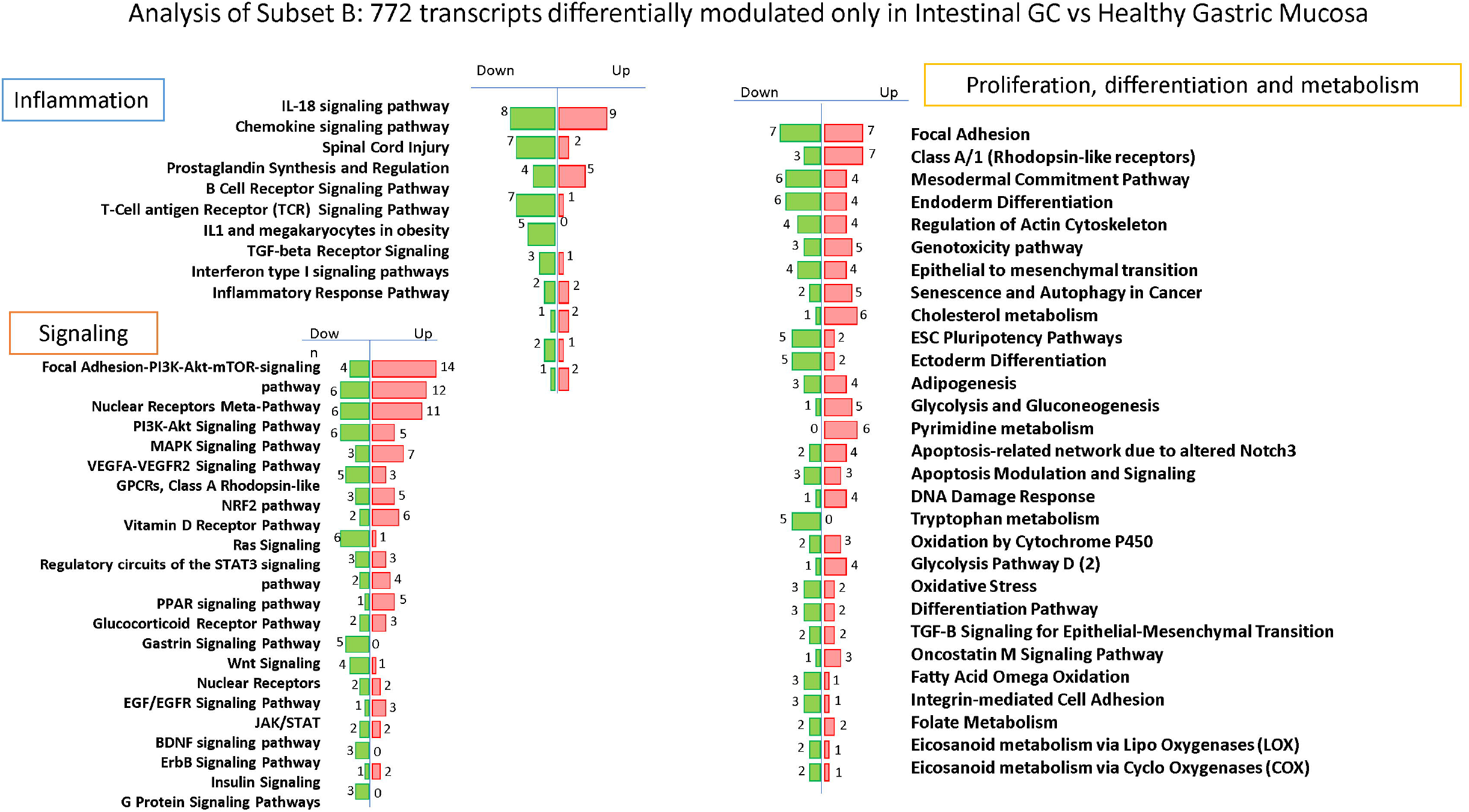
*Per* pathways analysis of Subset B. Analysis of 772 transcripts differentially modulated only in Intestinal Gastric Cancer vs Healthy Gastric Mucosa: identification of several pathways that can be grouped in three principal clusters: Proliferation, differentiation and metabolism, Inflammation and Signalling.

**Table 3.** Principal pathways of Inflammation Cluster for the subset B.

| Inflammation | Upregulated genes | Downregulated genes |
| --- | --- | --- |
| IL-18 signaling pathway | NCF2, <u>CCL4</u>, PLOD3, BID, HCAR2, <u>FN1</u>, <u>PTGS2</u>, TGM2, ULBP2 | FOXN3, KLF2, PRKCB, <u>CCL19</u>, DES, ACTA2, <u>NR0B2</u>, SPON1 |
| Chemokine signaling pathway | <u>CXCR2</u>, <u>CCL4</u> | <u>CCL19</u>, <u>CCL11</u>, <u>CXCL14</u>, VAV3, PRKCB, PRKACB, AKT3 |
| Spinal Cord Injury | E2F1, RTN4R, LILRB3, <u>PTGS2</u>, GJA1 | SLIT2, SLIT3, RGMA, VIM |
| Prostaglandin Synthesis and Regulation | <u>PTGS2</u> | PTGFR, PTGER1, HPGD, HPGDS, AKR1C3, AKR1C1, AKR1C2 |
| B Cell Receptor Signaling Pathway |  | CD79B, CD79A, BLNK, BLK, CR2 |
| T-Cell antigen Receptor (TCR) Signaling Pathway | RIPK2 | GRAP2, VAV3, VIM |
| IL1 and megakaryocytes in obesity | <u>TIMP1</u>, <u>S100A9</u> | SELENBP1, FCER1A |
| TGF-beta Receptor Signaling | BAMBI, <u>SERPINE1</u> | BMP4 |
| Interferon type I signaling pathways | SOCS3 | PDCD4, PTPRC |
| Inflammatory Response Pathway | <u>FN1</u>, <u>LAMA5</u> | CD40LG |

Among inflammatory pathways, we found also an overexpression of several genes identified as diagnostic or prognostic markers of gastric cancer including E2F1, that induces upregulation of lncRNA HCG18 thus stimulating proliferation and migration of gastric cancer [96], TIMP1, a key gene in the development of gastric cancer recognized as a potential prognostic marker when coexpressed with MMP-7 [97,98], S100A9, a diagnostic and prognostic biomarker in gastric cancer [99,100], and SERPINE1, highly expressed and significantly related to a poor prognosis of gastric adenocarcinoma [90]. Interestingly, also LAMA5 that promotes colorectal liver metastasis growth, resulted upregulated in intestinal gastric cancer [101].

The *per pathway* analysis of Signaling cluster in intestinal gastric cancer revealed a higher modulation of several signaling pathways such as PI3K-Akt-mTOR, MAPK, Ras, Jak/Stat, NFkB, VEGF (Table 4).

**Table 4.**
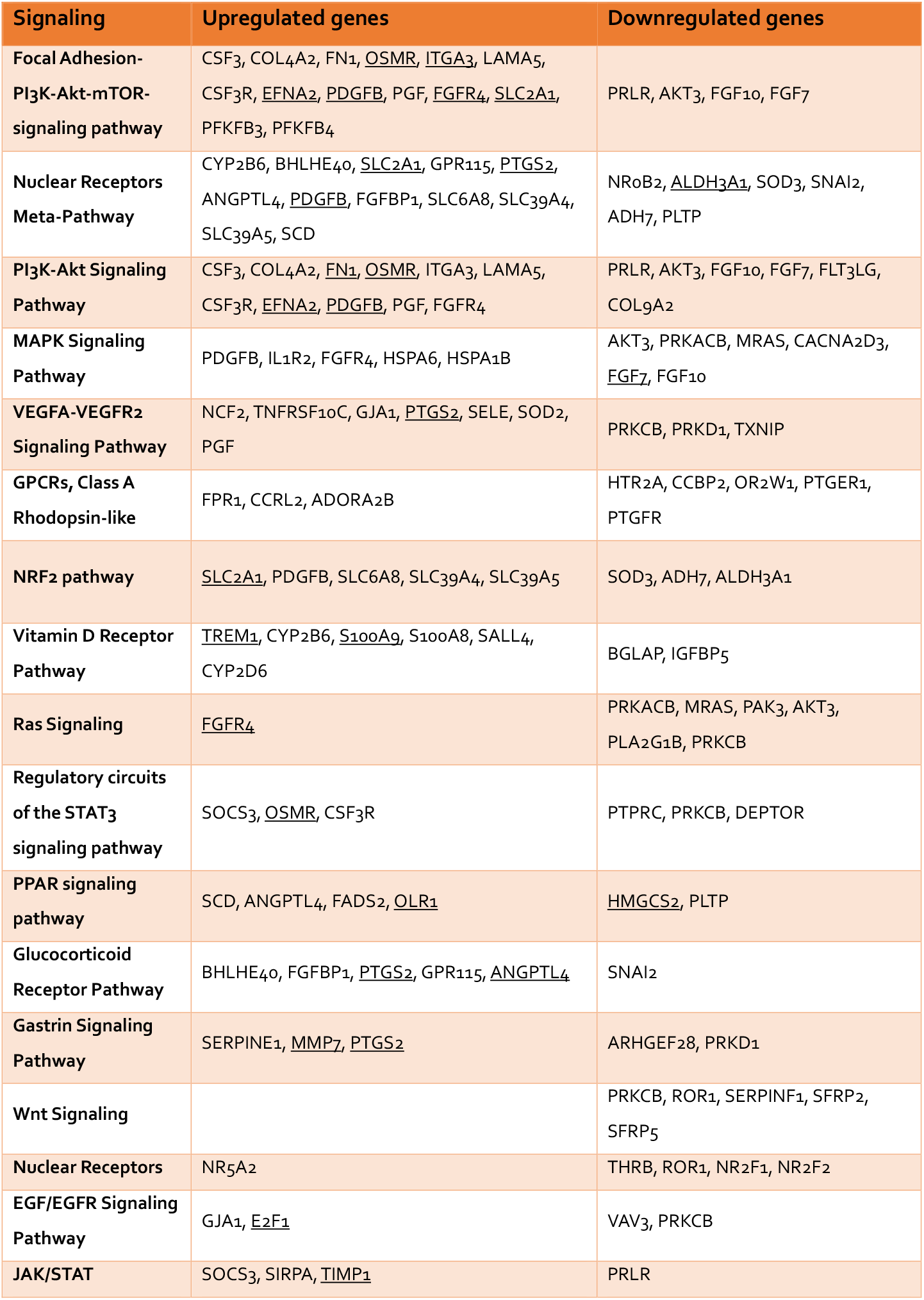
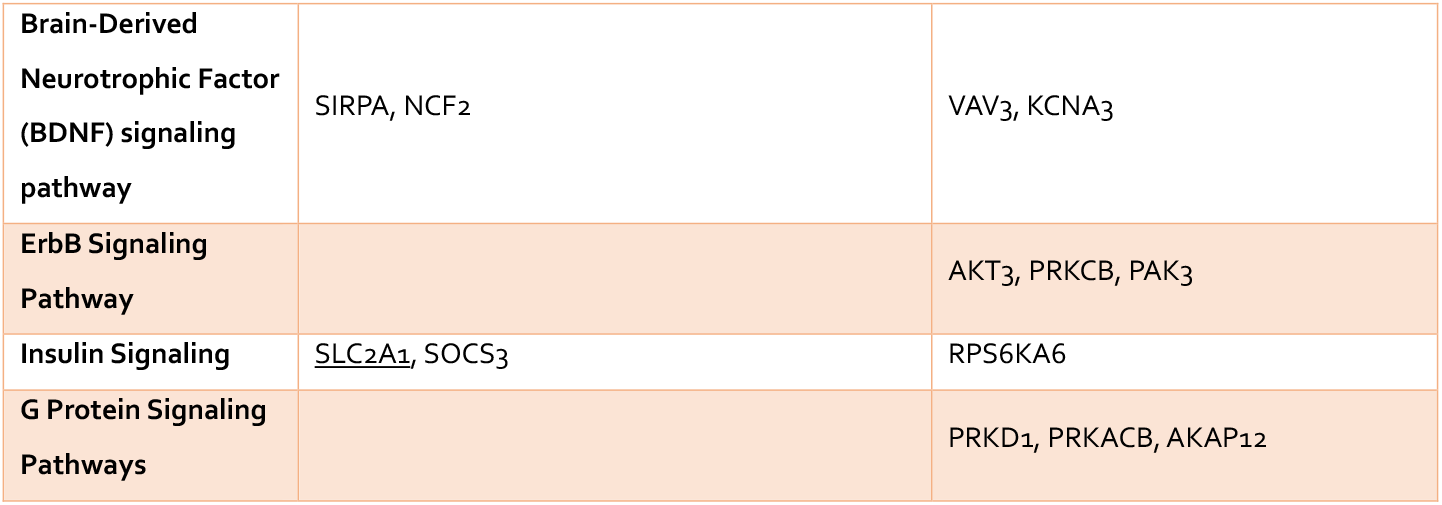
Principal pathways of Signaling Cluster for the subset B.

The pathway most modulated resulted the Focal Adhesion-PI3K-Akt-mTOR-signaling pathway with 14 upregulated genes and 4 downregulated genes. In particular we found an overexpression of OSMR, the receptor of Oncostatin M which promotes gastric cancer growth and metastasis [102], ITGA3, which promotes peritoneal metastasis and correlates with poor prognosis in patients with gastric cancer [103], EFNA2, one of the members of the ephrin family that are target of WNT/beta-catenin signaling implicated in the development of carcinogenesis [104], PDGFB, whose overexpression increases the growth, invasion, and angiogenesis of gastric carcinoma cells [105], FGFR4, that regulates proliferation and anti-apoptosis during gastric cancer progression [106], and SLC2A1, that induces tumor cell proliferation and metastasis in gastric Cancer when overexpressed [107].

Interestingly, among the other pathways we found an increased expression of OLR1, that facilitates metastasis of gastric cancer through driving EMT and PI3K/Akt/GSK3β activation [108], and MMP7, identified as prognostic marker in gastric cancer when co-expressed with TIMP1 [98]. Conversely HMGCS2, identified as tumor suppressor with prognostic impact in prostate cancer, resulted downregulated also in intestinal gastric cancer subset [109].

The analysis of Proliferation, differentiation and metabolism Cluster (Supplementary Table 6) revealed among others, the overexpression of several genes involved in invasion and metastasis of gastric cancer including CXCR1, that promotes malignant behavior of gastric cancer cells in vitro and in vivo inducing AKT and ERK1/2 phosphorylation [110,111], FPR2, that induces invasion and metastasis of gastric cancer cells and predicts the prognosis of patients acting as a novel prognostic marker [112], CARD14, involved in the progression from normal gastric epithelial to gastric cancer [113]. Finally, we found also a decreased expression of CLDN11, whose silencing is associated with increased invasiveness of gastric cancer cells [114]. Furthermore, the most upregulated genes specific for intestinal gastric cancer subset resulted OLFM4, involved in early gastric carcinogenesis and associated with prognostic significance in advanced stages [115], CDH17 and TFF3, specific markers of intestinal metaplasia gastric cancer patients [116], TRIM29, identified as oncogene in gastric cancer marker of lymph node metastasis [117,118], and SYT8, a promising target for the detection, prediction, and treatment of peritoneal metastasis of gastric cancer [119]. Conversely, among the transcripts more downregulated we found PSCA, whose reduced expression promotes gastric cancer proliferation and is related to poor prognosis [120], GKN2, which results downregulated in gastric cancer and when restored suppresses gastric tumorigenesis and cancer metastasis [121], and ALDH1A3, MLF1 and GREM1, all methylated at promoter level in gastric cancer [122].

### Functional enrichment analysis of diffuse gastric-specific genes and intestinal-specific genes

Finally, functional enrichment analysis was performed separately for gastric cancer histotype-specific genes, using DAVID tools [24] and the results were shown in Figure 5. For histotype-specific genes in diffuse gastric cancer tissues, functions including methylation, lipid metabolism (VLDL, Lipoprotein, Lipid Transport, Lipid metabolism, HDL), cell division and adhesion were significantly enriched (Figure 5). Conversely, the histotype-specific genes in intestinal gastric cancer samples were dramatically enriched in functions, including cell migration (cell membrane, extracellular matrix, metalloproteases, chemotaxis), vasculature development (angiogenesis, cell adhesion), apoptosis, but especially immune system regulation and inflammation (immunoglobulin domain, cytokine, innate immunity, cytokine-cytokine receptors interaction, adaptative immunity, prostaglandin metabolism, B cell activation) (Figure 5).

**Figure 5.**
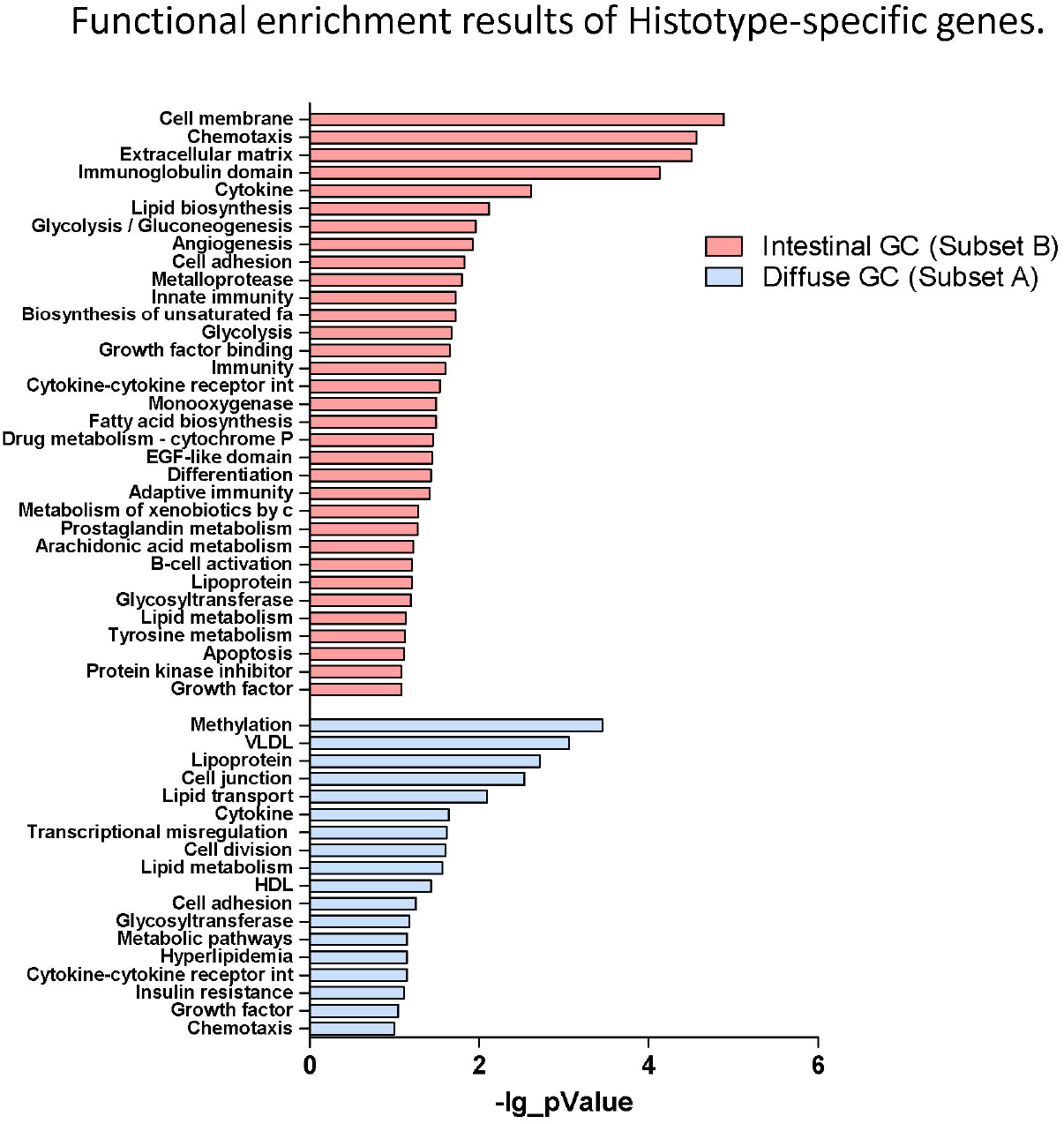
Functional enrichment analysis of Histotype-specific genes. Functional enrichment results of diffuse gastric-specific genes and intestinal-specific genes performed separately using DAVID tools.

Accordingly, these data confirmed that as described above intestinal histotype-specific genes were significantly enriched in biological processes such chemotaxis, inflammation, Innate and adaptative immunity, cellular adhesion, angiogenesis and modulation of several signaling pathways, such as PI3K-Akt-mTOR, MAPK, Erk1/2, Ras, Jak/Stat, NFkB, VEGF, involved in their regulation.

### Identification of histotype-specific genes

In order to refine the analysis, the selection criteria were strengthened with a threshold of FDR ≤0.1 and fold-change ≥3 applied. The stringent criteria generated a list of 7 upregulated and 0 downregulated transcripts in diffuse gastric cancer compared with healthy mucosa, whereas we found 14 upregulated transcripts and 11 downregulated transcripts in intestinal gastric cancer that met these criteria (Figure 6A and B).

**Figure 6.**
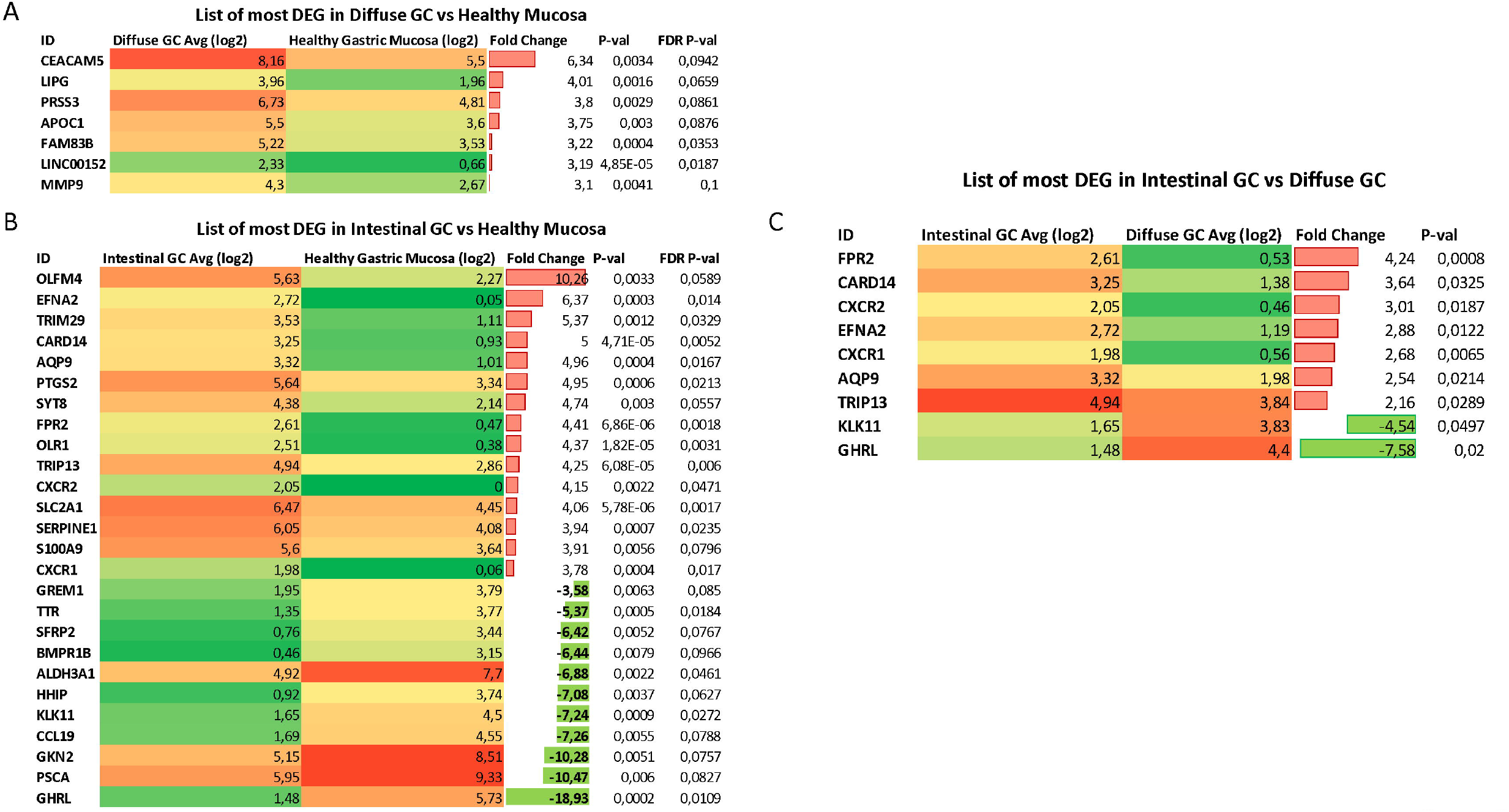
Identification of Histotype-specific genes. List of Differential Expressed Genes (DEGs) obtained using stringent criteria (threshold of FDR ≤0.1 and fold-change ≥3 applied) in (**A**) diffuse gastric cancer and (**B**) intestinal gastric cancer compared with healthy mucosa. (**C**) List of DEGs selected by using the stringent criteria between Intestinal gastric cancer and Diffuse gastric cancer samples.

Interestingly, we found 9 of these genes selected by using the stringent criteria, when we analyzed the Differential Expressed Genes (DEGs) between Intestinal gastric cancer and Diffuse gastric cancer samples (Figure 6C). In particular, genes encoding for N-formyl peptide receptor 2 (FPR2), Caspase recruitment domain-containing protein 1 (CARD14), C-X-C Motif Chemokine Receptor 2 (CXCR2), Ephrin A2 (EFNA2), C-X-C Motif Chemokine Receptor 1 (CXCR1), Aquaporin-9 (AQP9) and Thyroid Hormone Receptor Interactor 13 (TRIP13) resulted overexpressed in Intestinal Gastric cancer. Conversely, the expression of genes encoding for Ghrelin (GHRL) and Kallikrein Related Peptidase 11 (KLK11) was decreased in intestinal gastric cancer compared with diffuse histotype (Figure 6C). For these reasons, our results suggest that FPR2, CARD14, CXCR2, EFNA2, CXCR1, AQP9 TRIP13, along with GHRL and KLK11, could be used as promising biomarkers for gastric cancer diagnosis or prognostic evaluation.

### Clinical results

The 4-year overall survival (4Y-OS) of the study population is 52 % (Figure 7A). The Kaplan-Meier survival curve showed that intestinal group presented a 4Y-0S of 80%. Conversely, diffuse group has revealed a worse prognosis with only 25% of patients alive after four years from surgery resection (Figure 7B). The difference was statistically significant.

**Figure 7.**
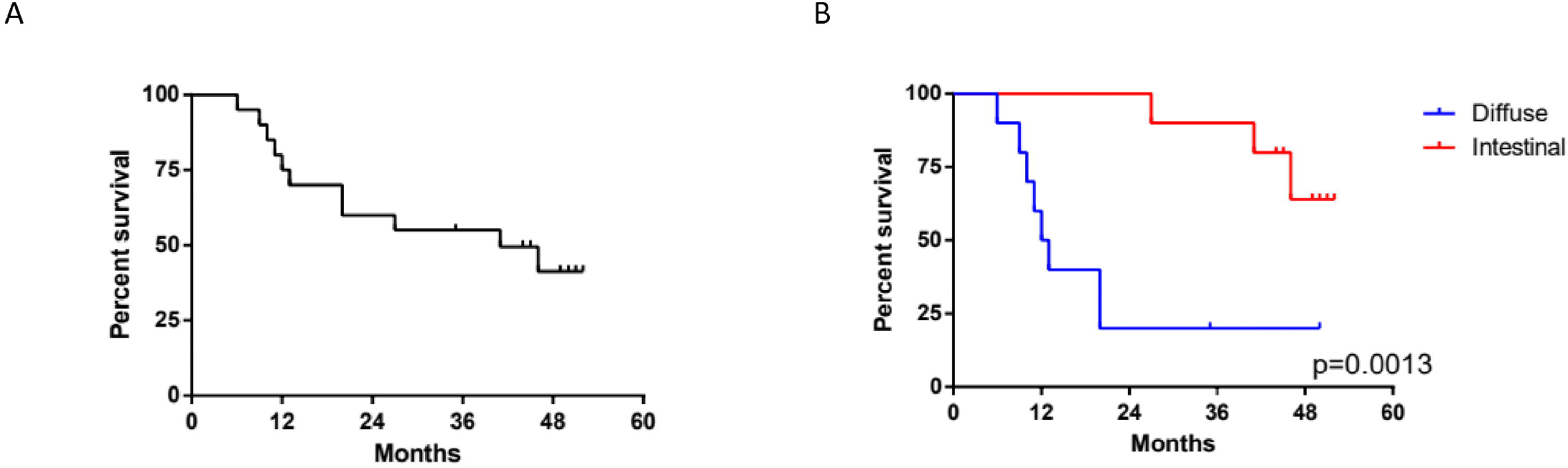
Clinical results. **(A)** 4-year overall survival (4Y-OS) of the study population. (**B**) Kaplan-Meier survival curve showing the 4Y-OS of intestinal and diffuse group. (p < 0.05).

According to the univariate analysis, performed as described in material and method section, the Intestinal group there was no clinical factor impacting the overall survival, whereas in the diffuse group, male sex and High N/L ratio worsen patients’ prognosis (Figure 8). In particular, in diffuse sub set females seems to have a better survival (4Y-OS: 40%) compared to males (4Y-OS: 0%) (Figure 8A). According to the median value of N/L ratio, patients were dichotomized into two subgroups. In the diffuse type, the survival was lower in the N/L higher group (40% vs 0%), as shown in figure 8C, in a statistically significant way. This was not confirmed in the intestinal subgroup (Figure 8D). In the diffuse group we eventually considered sex and N/L ratio status in multivariate analysis, but none of these factors turned out to be statistically independent.

**Figure 8.**
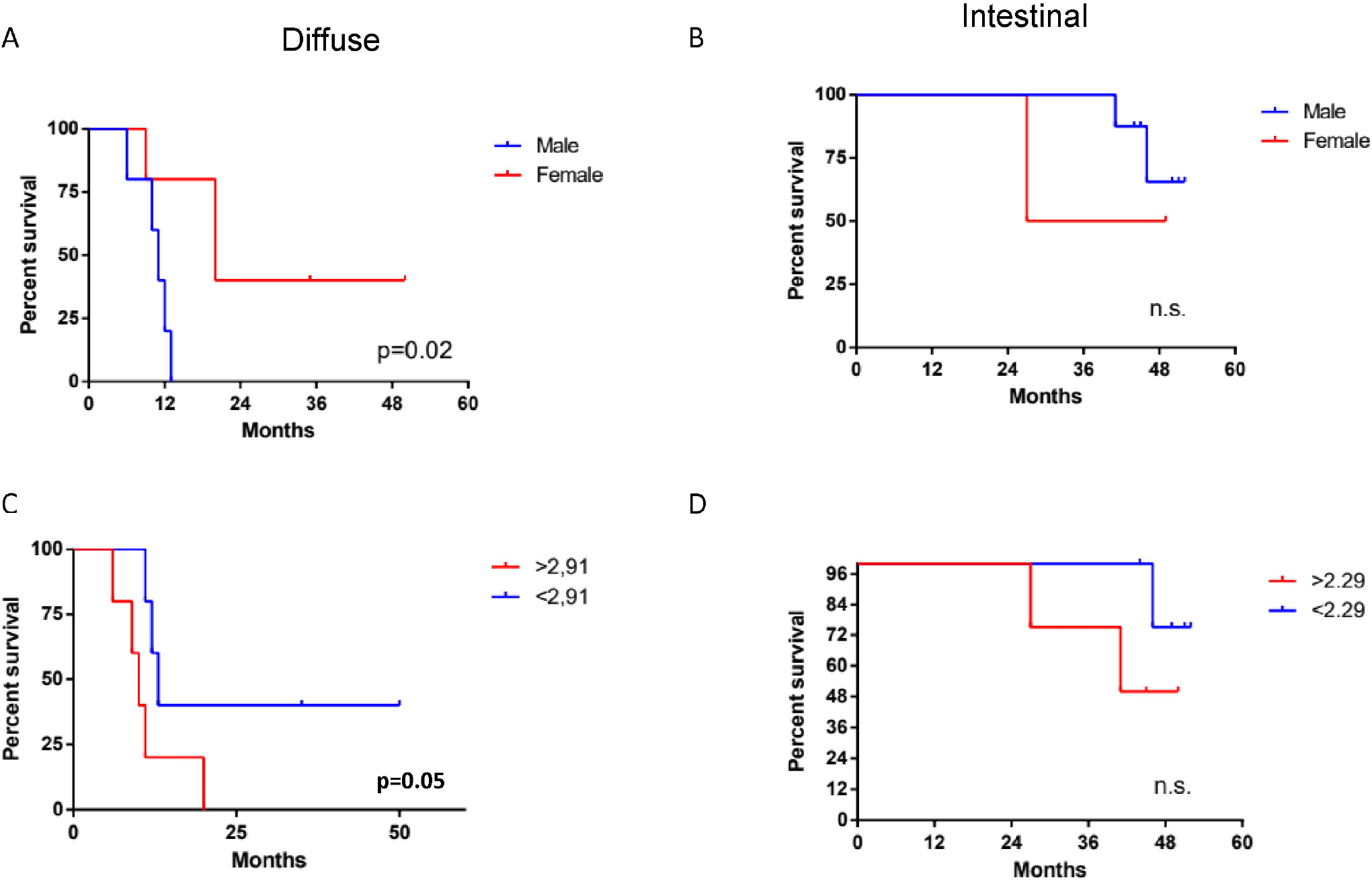
Univariate analysis. Analyisis of 4Y-OS performed as described in material and method section according sex (**panels A and B**) or N/L ratio status (**panels C and D**) in diffuse and intestinal groups. (p < 0.05).

At the end of our analysis we evaluated the impact of genes’ expression in overall survival. For this purpose, patients were dichotomized into 2 groups according to the median value of expression each gene (Figure 9A). In the intestinal group only AQP9 gene expression showed an impact in patient survival: patients having a high AQP9 expression show a significant worse prognosis compared to patients with low expression (31.5% vs 100% 4Y-OS). In the diffuse group the expression of two genes, CARD14 and CXCR2, revealed an impact in patient survival. Despite the lower expression of these genes in diffuse group compared to intestinal group, the higher expression of CARD14 or CXCR2 was significantly associated with a worse prognosis in patients of diffuse subset (Figure 9C ad D). In particular, patients with a higher expression of CARD14 or CXCR2 show a 4Y-OS of 0% compared to patients having a lower expression of these genes, in which the 4Y-OS was of 42% and 40% respectively (Figure 9C and D). Furthermore, we considered both CARD14 and CXCR2 expression in multivariate analysis, and we found that in particular the CXCR2 expression resulted a statistically independent factor (Table 5)

**Figure 9.**
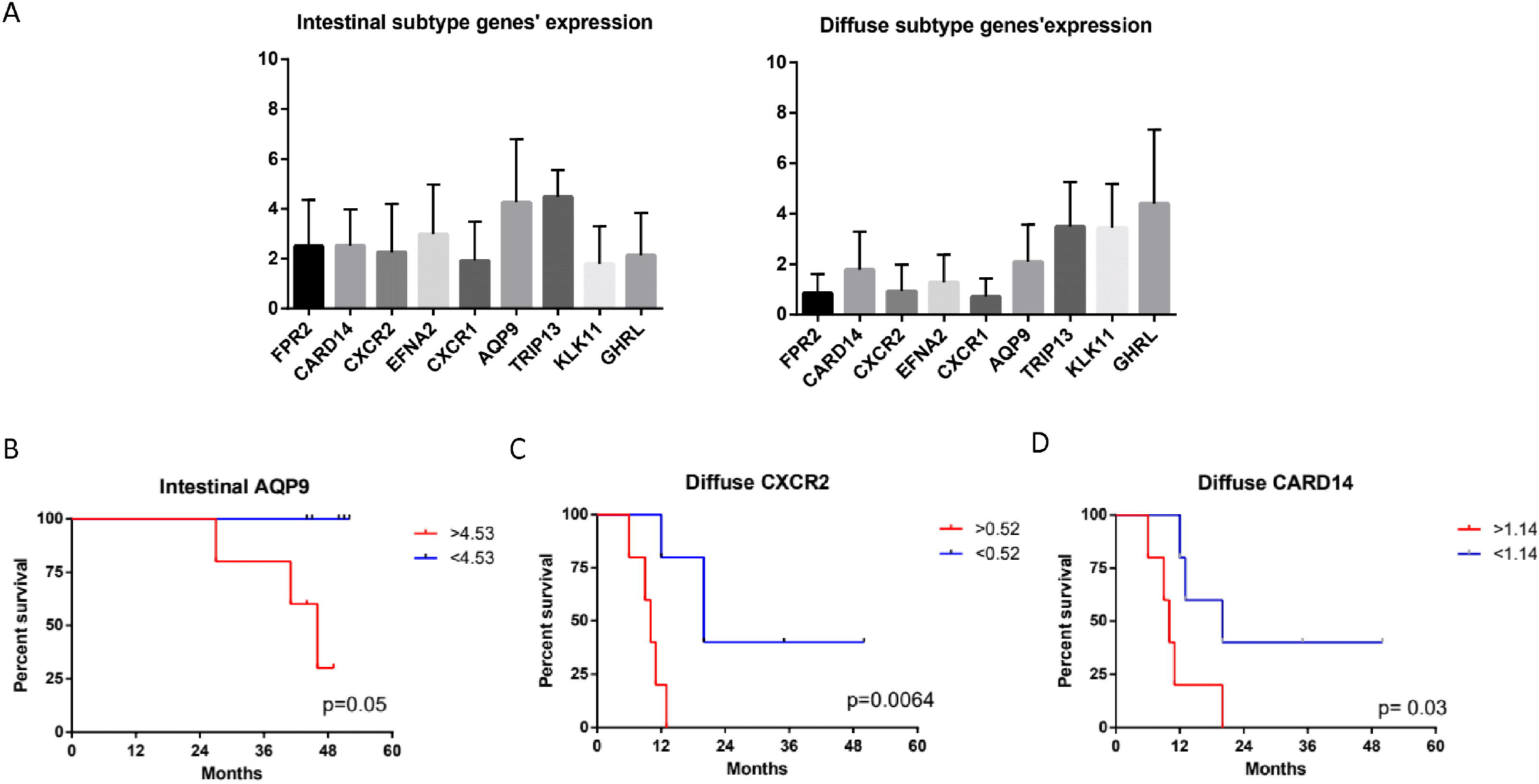
Impact of gene expression in overall survival. (**A**) Patients were dichotomized into 2 groups according to the median value of expression each gene. (B) Impact of AQP9 gene expression on patient survival in the intestinal group. Impact of (C) CXCR2 and (D) CARD14 on patient survival in the diffuse group. (p < 0.05).

**Table 5.** Multivariate analysis of gene expression for the diffuse sub set.

| Covariate | Regression<br>Coef. | SE | Regression<br>coef./SE | P | RR | 95% CI RR |
| --- | --- | --- | --- | --- | --- | --- |
| CARD 14 | 1,3882 | 0,9045 | 2,3556 | 0,128 | 4,0075 | 0,6869-23,3795 |
| CXCR2 | 2,5435 | 1,2094 | 4,4228 | 0,0355 | 12,7244 | 1,2033-134,5504 |

## Discussion

In the present study we report an integrative approach combining the transcriptome sequencing (RNA-seq) and clinical-pathological phenotypes to a series of gastric cancer recorded in a single center in Italy [123]. In the clinical setting, the histopathological classifications by Lauren and the WHO, presently remain the two classifications most commonly used for the therapeutic decisional process [9]. According to the Lauren classification, gastric cancers are subdivided into two major histological subtypes, namely intestinal type and diffuse type adenocarcinoma. Both the intestinal and the diffuse types have been associated with chronic gastritis and *H. pylori* infection, that represents the main cause of gastric cancer, however, the histological changes leading to intestinal type are better characterized [124], suggesting that the later could be considered the end-result of an inflammatory multistep process that starts with *H. pylori infection* and progresses to chronic gastritis, atrophic gastritis and finally to intestinal metaplasia and dysplasia [7,124]. Conversely, the a development cascade for diffuse gastric cancer type, which convey a worst prognosis, is less defined. Despite the Lauren classification has been proposed more than half a century ago it maintains several advantages in term of easy handling and prognostic significance [125]. Therefore, a more detailed knowledge of the molecular subsets of two pathology entities described by Lauren may lead to a newer approach to gastric cancer tailored treatment [126].

In this study we provide the results of in deep characterization of the transcriptome patterns from 12 diffuse and 12 intestinal gastric cancer patients using high-throughput sequencing technology. We identified 885 transcripts differentially expressed in comparison to non-neoplastic tissue by both the diffuse and intestinal gastric cancer, that were considered to represent a group of recurrently deregulated genes potentially associated with tumorigenesis. As described in result section we found in this subset of transcripts a modulation of cell cycle regulators including, several genes involved in Epithelial to Mesenchymal Transition, chemokine and cytokine signaling pathways with an upregulation of genes such as CCL3, CCL15 [43], CCL20 [44], CXCLs family [45,46] [47–50], IL1b, IL11 and IL8 [51–53].

Surprisingly, the analysis of diffuse gastric cancer subset revealed in addition to a modulation of immune and proliferation pathways, also an upregulation of genes involved in proliferation and lipid metabolism such as HNF4A [65], APOC1, [66], APOE, [67,68], FASN [69], while the expression of VLDLR [70] and CCL2 [72], were downregulated.

The analysis of intestinal gastric cancer subset show a strong inflammatory component, likely due to its close association with *H. pylori* infection, with a robust modulation of genes involved in chemotaxis, inflammation and innate and adaptive immunity. Moreover, the *per pathway* analysis revealed a higher modulation of several signaling pathways including PI3K-Akt-mTOR, MAPK, Ras, Jak/Stat, NFkB, VEGF.

These data were confirmed also by the functional enrichment analysis performed using DAVID tool. We found that the diffuse gastric cancer tissues were enriched for functions including methylation, lipid metabolism cell division and adhesion, whereas the intestinal gastric cancer group was dramatically enriched for genes involved in cell migration, vasculature development apoptosis, immune system regulation and inflammation.

The clinical analysis of our study population first confirmed that patients affected by diffuse type adenocarcinoma showed a worse prognosis with only 25% of patients survival after four years from surgical resection. None of the clinical factors investigated (Table 1) had any impact on intestinal group prognosis, while gender and preoperative inflammation status impacted on the survival of patients with diffuse gastric cancers, although at the multivariate analyses non of the clinical parameters maintained a statistical significance.

To better characterize the diffuse and intestinal gastric cancer transcriptome profile, we have strengthened the selection criteria for the analysis of differentially modulated genes (DEGs). Therefore, we have selected 9 genes differentially modulated between the intestinal and diffuse gastric cancer samples: FPR2, CARD14, CXCR2, EFNA2, CXCR1, AQP9, TRIP13, GHRL and KLK11, to be used as differentially expressed biomarkers for gastric cancers. In following this approach, we have first investigated whether the nine selected genes impact on overall patients survival. The results of this subset analysis demonstrated that in the intestinal group only the expression of AQP9 gene exerted a significant impact in patient survival. AQP9, a member of the aquaporin family, is involved in development of several tumors, promoting the proliferation, migration and invasion of tumor cells [127]. Previous studies have shown that AQP9 induces the growth and the migration of prostate cancer [128] and astrocytoma cells [129]. Conversely, AQP9 might inhibit the invasion of liver cancer cells and the proliferation of xenograft tumors [130], and also activates RAS signal and sensitize tumor cells to chemotherapy in colorectal cancer [131]. AQP9 had significant association with various immune infiltrating cells including CD8+ and CD4+ T cells, neutrophils, tumor associated macrophages (TAMs) and dendritic cells (DCs) [132,133]. However, high AQP9 expression was significantly correlated with worse prognosis in breast [134], colon and lung cancers [135], while predicted better prognosis in gastric cancer both in diffuse and intestinal gastric cancer [136]. Therefore, we can suppose that our results showing a worse prognosis for patients with high levels of AQP9 in intestinal gastric cancer subset, may be due to the presence of a stronger immune and inflammatory component. Moreover, it was been shown that gene markers of M2 macrophages were moderately to very strongly correlated with AQP9 expression [137], suggesting that AQP9 might be involved in the polarization of TAMs and in the immunosuppression in cancer. In the diffuse gastric cancer group, both CARD14 and CXCR2 expression, revealed an impact in patient survival. In this subset, the higher expression of CARD14 or CXCR2 was significantly associated with a worse patient prognosis. Importantly, the multivariate analysis revealed that the CXCR2 expression resulted a statistically independent factor.

CARD14, is a member of the Caspase recruitment domain family of proteins, which play an important role in immune and inflammatory response, and cell survival and proliferation. It is strongly expressed in the epidermal keratinocyte of the skin and is involved in inflammatory disorders of the human skin, such as psoriasis [138,139]. CARD14 isoforms are expressed in several hematopoietic cells and tissues such as bronchus, cervix, colon and lung as well as cancer cell lines derived from these tissues [140,141]. In particular, it was shown that CARD14 resulted overexpressed in breast cancer cell lines, and its knockdown led to decreased breast cancer cell proliferation and migration ability, accompanied by the induction of cell death through NF-κB [142].

CXCR2, is a potent pro-tumorigenic chemokine receptor that can induce inflammation in the tumor microenvironment by mediating the recruitment of different stromal cells and thus promoting the progression of cancer cells [143]. CXCR2-targeted therapy has shown previously promising results in several solid tumors, including breast cancer [144], pancreatic cancer [145], and rhabdomyosarcoma [146]. More recently, its expression has been associated with the prognosis of patients with gastric cancer [48]. CXCR2 ligands, produced by TAM, significantly promote proliferation and migration of gastric cancer cells through activating a CXCR2/STAT3 feed-forward loop [147]. Gastric cancer cells in turn, secrete TNF-α to induce the release of CXCR2 ligands from macrophages [48]. Furthermore, CXCR2 along with CXCR4 overexpression, was associated with more advanced tumor stages and poorer survival in gastric cancer patients [94]. CXCR4 and CXCR2 activate NF-κB and STAT3 signaling, while NF-κBp65 can then transcriptionally activate CXCR4 and STAT3 can activate CXCR2 expression [94]. It was been shown that this crosstalk between CXCR4 and CXCR2 contributed to EMT, migration and invasion of gastric cancer. Conversely, the inhibition of CXCR2 pathway in gastric cancer cells suppressed migration and metastasis of gastric cancer both in vitro and in vivo [48]. Therefore, the interaction between CXCR2 and tumor microenvironment results of critical importance for tumor progression [148]. Recent studies have been demonstrated that the blockade of the receptor with specific inhibitors [149], as well as the inhibition of the recruitment of immune cells via the CXCL1-CXCR2 axis [143], appear a promising therapy for gastric cancer primarily for diffuse subtype.

In summary our analysis detected 7 genes up regulated in the Intestinal type and 2 genes down regulated when compared to the healthy mucosa, with AQP9 expression influencing also patients prognosis. In the diffuse type, with an *ab-initio* worse prognosis, we were able to detect two genes, CARD14 and CXCR2, impacting prognosis. In particular CXCR2 seems to play a key role, resulting the only factor independently impacting overall survival.

The present study suggests that targeting AQP9 and CXCR2 may represent a novel strategy for gastric cancer therapy, in intestinal and diffuse patients respectively. However, further studies will be needed to confirm the role of these genes as therapeutic targets and biomarkers in gastric cancer.

## Acknowledgment

This work was supported by a grant from Regione Campania-POR Campania FESR 2014/2020 ***“Combattere la resistenza tumorale: piattaforma integrata multidisciplinare per un approccio tecnologico innovativo alle oncoterapie-Campania Oncoterapie” (Project N.* B61G18000470007)**

