## Supplementary Tables for "Analysis of gastric cancer transcriptome allows the identification of histotype specific molecular signatures with prognostic potential"

| Proliferation, Differentiation and Metabolism | Up-regulated genes | Down-regulated genes |
|---|---|---|
| **Cell Cycle** | PLK1, PCNA, ESPL1, MCM2, CDC25B, CCNA2, CDC45, PTTG1, CDK1, PKMYT1, CDC20, CCNE1, CDC25C, SFN, CDC6, MCM3, MCM4, ORC1, ORC6, BUB1, CCNB2, CCNB1, E2F5, E2F2, E2F3, CDC25A, TTK | CDKN1C |
| **Mitotic G2-G2/M phases** | CDC25C, MYBL2, TPX2, PKMYT1, AJUBA, CCNB1, CDC25B, CDC25A, PLK1, BORA, FOXM1, CCNB2, CDK1, AURKA, CENPF, GTSE1, HMMR | |
| **Ectoderm Differentiation** | TNFRSF11B, HIST1H2BH, RHPN1, LY6E, TFAP2A | PLCXD3, FZD4, LDB2, ARHGDIG, PRKAG2, ARHGAP15, GLI3, ZBTB16, PTPN13, ARHGAP10 |
| **Mitotic G1-G1/S phases** | MCM10, TOP2A, CDC25A, CDC45, CCNE1, CCNA2, RRM2, CDK1, MYBL2, DHFR, CDC6, ORC1, TK1, TYMS, PCNA | |
| **Senescence and Autophagy in Cancer** | PLAU, CDC25B, CXCL1, IL8, PCNA, IL1A, IL1B, CCL3, TNFSF15, INHBA, HMGA1 | IGF1, BCL2, GABARAPL1 |
| **Epithelial to mesenchymal transition (EMT)** | GDF15, TMPRSS4, CLDN1, COL4A1, FOXM1, CLDN3, CLDN4, CLDN7, WNT5A | EIF5A2, COL4A3, COL4A5, WNT2B, FZD4 |
| **Focal Adhesion** | ITGA2, SPP1, LAMC2, COL4A1 | LAMA2, PDGFD, CHAD, BCL2, ITGA8, COL2A1, IGF1, TNXB, MAPK10 |
| **Circadian rhythm related genes** | TYMS, CLDN4, TNFRSF11A, TOP2A | ADA, SLC9A3, NTRK3, ID4, PRKAA2, PTGDS, RORB, MAPK10 |
| **G1 to S cell cycle control** | E2F2, PCNA, CDK1, CCNE1, CCNB1, MCM4, MCM3, MCM2, E2F3, CDC25A | CDKN1C |
| **Metapathway biotransformation Phase I and II** | HS3ST1 | CYP4B1, GSTA1, GLYATL1, GPX3, CYP1B1, SULT2A1, GSTM2, GSTM5, FMO2 |
| **Adipogenesis** | LIF, HMGA1, OSM, AGPAT2 | IGF1, PTGIS, SLC2A4, LIFR, RXRG, KLF15 |

| | | |
|---|---|---|
| **DNA Damage Response** | PMAIP1, FANCD2, CDC25A, SFN, CCNB1, CDC25C, CCNE1, CCNB2, RAD51 | |
| **Regulation of Actin Cytoskeleton** | VIL1, F2R | CHRM3, FGF14, PIK3C2G, FGF2, CYFIP2 |
| **DNA Replication** | RFC4, PCNA, MCM2, MCM10, MCM4, CDC6, MCM3 | |
| **ESC Pluripotency Pathways** | WNT5A, LIF | FZD4, LIFR, FGF14, FGF2, WNT2B |
| **Mesodermal Commitment Pathway** | INHBA, WDHD1, NCAPG2 | PBX3, SLC2A12, TOX, FZD4 |
| **Endoderm Differentiation** | WDHD1, NCAPG2 | SFRP1, TCEAL2, PBX3, SLC2A12, TOX |
| **Differentiation Pathway** | INHBA, IL11, WNT5A | FGF2, KIT, IGF1, WNT2B |
| **Cell Cycle Checkpoints** | BUB1B, CDC20, CDC25A, CDC25C, PKMYT1, CLSPN, MAD2L1 | |
| **Prostaglandin Synthesis and Regulation** | S100A10, ANXA2S, OX9 | PTGIS, PTGDS, PTGER3 |
| **Glycolysis and Gluconeogenesis** | PFKP, LDHA, ENO1, HK2 | SLC2A4, FBP2 |
| **Synthesis of DNA** | MCM3, CDC45, FEN1, CDC6, GINS1, MCM2 | |
| **Integrin-mediated Cell Adhesion** | ITGA2 | SEPP1, CAPN6, ITGA8, ITGAL, MAPK10 |
| **Trans-sulfuration pathway** | LDHA | CKM, CDO1, CKB, CKMT2 |
| **M/G1 Transition** | ORC1, MCM10, CDC6, CDC45, ORC6 | |
| **Apoptosis** | PMAIP1, BIRC5 | MAPK10, IGF1, BCL2 |
| **Matrix Metalloproteinases** | MMP12, MMP10, MMP3, MMP1 | TIMP3 |
| **Oxidative Damage** | PCNA | CDKN1C, BCL2, MAPK10 |

**Supplementary Table 1. Principal pathways of Proliferation, differentiation and metabolism Cluster for the subset AB.**

| Inflammation | Up-regulated genes | Down-regulated genes |
|---|---|---|
| **IL-18 signaling pathway** | CCL3, MMP3, CLDN1, PLA2G7, IL1B, CCL20, CXCL16, MMP1, LMNB2, ENO1 | BCL2, SYT10, CD36 |
| **Chemokine signaling pathway** | CXCL3, CCL20, CCL3, CCL15, CXCL5, CXCL16, LYN | CCL21, CXCL12, GNG7, ADCY2, DOCK2 |
| **Spinal Cord Injury** | IL1B, MMP12, CXCL1, IL1A, CXCL2, E2F5, COL4A1, SOX9, CDK1 | AQP4, COL2A1 |
| **Breast cancer pathway** | RAD51, E2F2, E2F3, WNT5A | IGF1, KIT, PGR, FGF2, FZD4, WNT2B |
| **Complement and Coagulation Cascades** | F2R, PLAU, PLAUR | F10, C7 |
| **Cytokines and Inflammatory Response** | IL1A, IL1B, CXCL2, CXCL1, IL11 | |
| **Signal transduction through IL1R** | IRAK2, IL1RN, IL1A, IL1B | |
| **Cells involved in local acute inflammatory response** | IL1A, IL8 | C7, ITGAL |
| **LTF danger signal response pathway** | IL1A, IL1B, IL8 | LTF |
| **Interleukin-6 family signaling** | IL11, LIF, OSM | LIFR |
| **FasL pathway and Stress induction of HSP regulation** | IL1A, LMNB1, LMNB2 | BCL2 |
| **Interleukin-11 Signaling Pathway** | IL11, BIRC5, ITGA2 | BCL2 |

**Supplementary Table 2. Principal pathways of Inflammation Cluster for the subset AB.**

| Signaling | Up-regulated genes | Down-regulated genes |
|---|---|---|
| **PI3K-Akt signaling Pathway** | CCNE1, LAMC2, ITGA2, COL4A1, SPP1, ANGPT2, EFNA3, EPHA2, OSM, F2R, PIK3AP1 | CHAD, COL4A5, ITGA8, COL2A1, TNXB, LAMA2, COL4A3, GHR, ANGPT1, FGF14, FGF2, IGF1, PDGFD, KIT, LPAR1, GNG7, PRKAA2, PPP2R3A, BCL2 |
| **Focal Adhesion-PI3K-Akt-mTOR-signaling pathway** | LAMC2, ITGA2, COL4A1, SPP1, ANGPT2, EFNA3, EPHA2, OSM, F2R | CHAD, ITGA8, COL2A1, TNXB, LAMA2, ITGAL, GHR, ANGPT1, FGF14, FGF2, IGF1, PDGFD, KIT, LPAR1, GNG7, CAB39L, PRKAA2, PPP2R3A, SLC2A4, HIF3A |
| **Nuclear Receptors Meta-Pathway** | SLC7A11, EPHA2, CCL20, IL11, IL1B, MYOF, ENC1, TNS4, CDK1, SLC6A14 | SDPR, PDK4, TSC22D3, GPX3, SULT2A1, GSTA1, CYP1B1, SLC2A4, CDKN1C, PMP2, SLC2A12, DNER, GSTM5, GSTM2, SLC5A5 |
| **Vitamin D Receptor Pathway** | MXD1, KLK6, SPP1, S100A2, CD9, TNFRSF11B, CST1, CCNE1 | TRPV6, SULT2A1, ID4, SLC2A4, TIMP3, DNER, SFRP1, BMP6 |
| **NRF2 pathway** | SLC7A11, EPHA2, SLC6A14 | GSTA1, GPX3, SLC2A12, SLC2A4, GSTM5, GSTM2, SLC5A5 |
| **Gastrin Signaling Pathway** | BIRC5, ANXA2, IL8, CLDN1 | CHGA, HDC, CCKBR, SLC9A3, KIT |
| **Wnt Signaling** | PLAU, WNT5A, FOSL1 | WNT2B, FZD4, CAMK2B, MAPK10, SFRP1, PRICKLE2 |
| **VEGFA-VEGFR2 Signaling Pathway** | PLAU, SHB, TEAD4, MMP10, PLAUR | ITPR1, RKAA2, BCL2, TNXB |
| **Glucocorticoid Receptor Pathway** | IL11, ENC1, CCL20, TNS4 | PMP2, TSC22D3, CDKN1C, DNER, SDPR |
| **MAPK Signaling Pathway** | IL1A, CDC25B, IL1B | MAPK10, CACNA1A, CACNA2D2, FGF2, FGF14 |
| **Regulation of toll-like receptor signaling pathway** | CD80, IRAK2, SPP1, CCL3, PLK1, IL1B, IL8 | MAPK10 |
| **GPCRs, Class A Rhodopsin-like** | F2R | CHRM3, DRD5, HTR1E, CCKAR, CCKBR, PTGER3 |
| **Wnt Signaling Pathway and Pluripotency** | PLAU, WNT5A, FOSL1 | WNT2B, MAPK10, FZD4, PPP2R3A |

| | | |
|---|---|---|
| **PPAR signaling pathway** | SLC27A2, MMP1 | RXRG, FABP3, CD36, FABP4, ACADL |
| **G-Protein Signaling Pathways** | | GNG7, GNAZ, GNAO1, ADCY2, ITPR1, PRKAR2B, PDE1A |
| **JAK/STAT** | IL1RN, IL1B, AURKA | GHR, LEPR, IGF1, PRKAA2 |
| **Toll-like Receptor Signaling Pathway** | CD80, SPP1, CCL3, IL1B, IL8 | MAPK10 |
| **Ras Signaling** | EPHA2, CALML4, ETS2 | KIT, GNG7, MAPK10 |
| **TGF-beta Signaling Pathway** | CCNB2, CDK1, E2F5, MMP12, MMP1, ITGA2 | |
| **ErbB Signaling Pathway** | EREG | ERBB4, CAMK2B, MAPK10 |
| **TGF-beta Receptor Signaling** | SPP1, LIF, INHBA | ZNF423 |
| **Insulin Signaling** | | PIK3C2G, MAPK10, SLC2A4, PRKAA2 |
| **Regulatory circuits of the STAT3 signaling pathway** | F2R | GHR, LIFR, MAPK10 |
| **EGF/EGFR Signaling Pathway** | AURKA, MYBL2, PCNA | SH3GL2 |
| **Regulation of mitotic cell cycle** | NEK2, PTTG1, CDC20, BUB1B | |
| **Transcriptional regulation by the TAFP2** | MYBL2, ATAD2, NOP2 | KIT |
| **Leptin signaling pathway** | IL1RN, IL1B | LEPR, PRKAA2 |
| **Regulation of DNA replication** | CDC6, ORC1, MCM3, MCM2 | |
| **Cell surface interactions at the vascular wall** | EPCAM, ANGPT2 | ANGPT1, JAM2 |
| **GPCRs, Other** | F2R | GPR133, CHRM3, CCKBR |

**Supplementary Table 3. Principal pathways of Signaling Cluster for the subset AB.**

| Inflammation | Up-regulated genes | Down-regulated genes |
|---|---|---|
| **IL-18 signaling pathway** | IRAK1, IL18, IFNG, TOMM40, HMOX1, PYGB, ZC3H12A | CCL2, TRAF1, ACACB, TF, CA11 |
| **Spinal Cord Injury** | IFNG, MMP9, ANXA1, NOX4 | CCL2, PLA2G6, XYLT1 |
| **IL1 and megakaryocytes in obesity** | MMP9, IRAK1, IL18, IFNG | CCL2 |
| **Type II interferon signaling (IFNG)** | IFNG, PSMB9, OAS1 | |
| **Interleukin-6 family signaling** | CLCF1 | CRLF1, CNTFR |
| **Chemokine signaling pathway** | CCL22, CCL28 | SHC2 |
| **Development and heterogeneity of the ILC family** | IFNG, IL18, AREG | |
| **TNF alpha Signaling Pathway** | NOXO1 | TRAF1, CCL2 |
| **TNF related weak inducer of apoptosis Signaling Pathway** | MMP9 | TRAF1, CCL2 |
| **Prostaglandin Synthesis and Regulation** | ANXA4, ANXA1 | |
| **Toll-like Receptor Signaling Pathway** | IRAK1 | IFNA16 |

**Supplementary Table 4. Principal pathways of Inflammation Cluster for the subset A.**

| Signaling | Up-regulated genes | Down-regulated genes |
|---|---|---|
| **VEGFA-VEGFR2 Signaling Pathway** | SPHK1, ANXA1, RND1, F3 | ACACB, GAB1, SHC2, PRKCE, CCL2, ADAMTS1 |
| **Nuclear Receptors Meta-Pathway** | IFNG, FASN, GCLM, HMOX1, SLC6A20 | CCL2, FKBP5, GSTA4 |
| **Histone Modifications** | HIST1H3D, HIST1H3J, HIST1H3C, HIST1H3G, HIST1H4L | PRDM2 |
| **Insulin Signaling** | TRIB3, GRB14 | SHC2, GAB1 |
| **Ras Signaling** | RIN1 | SHC2, GAB1, PLA2G6 |
| **PI3K-Akt Signaling Pathway** | LAMB3, CREB3L1 | IFNA16, COL9A3 |
| **NRF2 pathway** | GCLM, HMOX1, SLC6A20 | GSTA4 |
| **Brain-Derived Neurotrophic Factor (BDNF) signaling pathway** | DPYSL2 | SHC2, ACACB, GRIA3 |
| **ErbB Signaling Pathway** | AREG | GAB1, SHC2 |
| **Statin Pathway** | APOC2, APOC1, APOE | |
| **G Protein Signaling Pathways** | | PDE1C, PDE8B, PRKCE |
| **Vitamin D Receptor Pathway** | SLC34A2, SLC37A2 | ADRB2 |

**Supplementary Table 5. Principal pathways of Signaling Cluster for the subset A.**

| Proliferation, Differentiation and Metabolism | Up-regulated genes | Down-regulated genes |
|---|---|---|
| **Focal Adhesion** | LAMA5, MYLK, COL4A2, FN1, PGF, ITGA3, PDGFB | PAK3, FIGF, BLK, AKT3, PRKCB, PARVG, VAV3 |
| **Class A/1 (Rhodopsin-like receptors)** | HCAR3, HCAR2, CXCR1, FPR1, FPR2, CXCR2, CCRL2 | PTGER1, PTGFR, CCL19 |
| **Mesodermal Commitment Pathway** | EXT1, SNAI1, BHLHE40, C1QBP | SESN1, CSRP2, BMP4, VAV3, SOX21, TET1 |
| **Endoderm Differentiation** | C1QBP, NME1, EXT1, EZH2 | SOX21, FOXN3, OTX2, TET1, SESN1, VAV3 |
| **Regulation of Actin Cytoskeleton** | FGFR4, MYLK, FN1, PDGFB | FGF10, MRAS, FGF7, PAK3 |
| **Genotoxicity pathway** | PLK3,E2F7,HIST1H2BN,IKBIP,DUSP14 | ARRDC4, ACTA2, TNFRSF17 |
| **Epithelial to mesenchymal transition in colorectal cancer** | SNAI1, FN1, EZH2, COL4A2 | AKT3, SNAI2, ZEB2, CLDN11 |
| **Senescence and Autophagy in Cancer** | E2F1, SERPINE1, FN1, CXCR2, SLC39A4 | CXCL14, IGFBP5 |
| **Cholesterol metabolism** | DHCR7, EBP, SQLE, ELOVL2, FADS2, SCD | HMGCS2 |
| **ESC Pluripotency Pathways** | PDGFB, FGFR4 | AKT3, FGF10, BMPR1B, FGF7, BMP4 |
| **Ectoderm Differentiation** | PODXL, ELOVL2 | DMD, CLDN11, RGMA, NR2F2, BMP4 |
| **Adipogenesis** | SERPINE1, SCD, E2F1, SOCS3 | BMP4, NR2F1, PRLR |
| **Glycolysis and Gluconeogenesis** | ALDOA, GAPDH, PGK1, SLC2A1, PGAM1 | ALDOB |
| **Pyrimidine metabolism** | CDA, TYMP, RRM1, NME1, PNP, POLR2H | |
| **Apoptosis-related network due to altered Notch3** | RIPK2, HELLS, CARD14, SOCS3 | VIM, VAV3 |

| | | |
|---|---|---|
| **Apoptosis Modulation and Signaling** | IL1R2, BAG3, BID | BLK, PRKD1, BMF |
| **DNA Damage Response** | BID, E2F1, CCNB3, H2AFX | SESN1 |
| **Tryptophan metabolism** | | ACAT1, INMT, ALDH1A1, HAAO, TPH1 |
| **Oxidation by Cytochrome P450** | CYP2B6, CYP2D6, CYP4F3 | CYP4X1, CYP4Z1 |
| **Glycolysis Pathway D (2)** | ALDOA, GAPDH, PGK1, PGAM1 | ALDOB |
| **Oxidative Stress** | SOD2, XDH | NFIX, SOD3, MAOA |
| **Differentiation Pathway** | CXCR1, PDGFB | BMP4, FGF10, FLT3LG |
| **TGF-B Signaling in Thyroid Cells for Epithelial-Mesenchymal Transition** | FN1, SNAI1 | VIM, SNAI2 |
| **Oncostatin M Signaling Pathway** | SOCS3, OSMR, SERPINE1 | PRKCB |
| **Fatty Acid Omega Oxidation** | CYP2D6 | ADH1C, ALDH1A1, ADH7 |
| **Integrin-mediated Cell Adhesion** | ITGA3 | VAV3, PAK3, AKT3 |
| **Folate Metabolism** | SLC19A1, SOD2 | FOLR4, SOD3 |
| **Eicosanoid metabolism via Lipo Oxygenases (LOX)** | FPR2 | HPGD, LTC4S |
| **Eicosanoid metabolism via Cyclo Oxygenases (COX)** | PTGS2 | HPGD, PTGFR |

**Supplementary Table 6. Principal pathways of Proliferation, differentiation and metabolism Cluster for the subset B.**
